## supplementary figures and suppl legends for "Optineurin links Hace1-dependent Rac ubiquitylation to integrin-mediated mechanotransduction to control bacterial invasion and cell division"

### LEGENDS TO SUPPLEMENTAL FIGURES

#### SUPPLEMENTAL FIGURE 1

**A)** Quantification of Ec bound to cells cultured on ECM of increasing stiffness (8, 25, 50 kPa). CNF1 was added at 1nM. Graph displays absolute number of CFU/ml (Y axis starts at $10^6$). Bars represent mean ± SEM of three independent experiments and 3 replicates per condition, MOI100. P values are calculated by one-way Anova with Dunnett's correction test for multiple comparison: non significant.

**B)** Quantification of phospho-protein levels for phospho-FAK (tyr-397), phospho-Paxillin (tyr-118), phospho-Src (tyr-416) and phospho-p130CAS (tyr-410) in HUVEC infected either with Ec or EcΔFimH, in presence of 1nM CNF1. Graphs display levels of phospho-proteins determined by immunoblotting relative to the total amount of GAPDH and normalized to the non-treated condition. Bars represent means ± SEM of three independent experiments. P values are calculated with two-way Anova with Tukey's correction test for multiple comparisons: ****, P ≤0.0001.

**C)** On the left: Quantification of internalized Ec in cells knocked down for beta1 integrin (siβ1) or control (siCTRL) after 30 minutes of gentamicin treatment in presence of 1nM CNF1. Graph displays absolute number of CFU/mL. Bars represent means ± SEM of three independent experiment and 3 replicates per condition. P values are calculated with one-way Anova: ****, P ≤0.0001. On the right: One representative immunoblot for knock-down verification of siβ1, GAPDH is used as a loading control.

**D)** Immunoblot quantification of phosphorylated FAK and SRC in siCTRL (white) and siβ1 (hatched) transfected cells, either non-treated (-) of upon Ec infection in presence of 1 nM CNF1 for 30 minutes. Data are normalized to siCtrl non treated condition. Mean ± SEM of three independent experiments. P values are calculated with two-way Anova: ***, P ≤0.001.

**E)** Representative Immunoblots showing knock down validation of Talin (TLN), Vinculin (VCL) and ICAP-1 in experiments quantified in Figure 1D. GAPDH was used as a loading control.

**F)** Quantification of cell-bound Ec to siCTRL, siVCL, siTLN or siICAP1 transfected cells in presence of 1nM CNF1. Graph displays the quantification of CFU/ml relative to the control condition (siCTRL). Mean ± SEM of three (TLN, VCL) or five (ICAP1) independent experiments. P values are calculated with one-way Anova: non significant.

**SUPPLEMENTAL FIGURE 2**

**A)** Principal component analysis visualization of the four replicates per condition. Different stiffnesses are represented by different colours, each replicate by a different dot shape.

**B)** Barplots of significant regulated proteins grouped for non treated (NT) and CNF1 treated conditions (CNF1). Y axis represents the number of proteins in the defined comparisons. A detailed description of the analysis can be found in the methods section.

**SUPPLEMENTAL FIGURE 3**

**A)** Immunoblots showing OPTN ubiquitylation profile. Cells were transfected with expression vectors for Histidine-tagged ubiquitin, HA-OPTN and Flag-HACE1 when indicated, with or without Myc-Rac1Q61L. Covalently bound His-ubiquitylated proteins were purified (HIS-P) and resolved on 12% SDS-PAGE before anti-HA immunoblotting to reveal ubiquitylated OPTN. Input corresponds to immunoblots on 0.5% total lysate to control protein expression levels.

**B)** Immunoblot showing levels of active Rac1 (Rac1-GTP) in HUVECs transfected with siCtrl or siOPTN and stimulated by 1h adhesion on fibronectin at different concentrations (Fn, 2.5, 7.5 and 15 µg/ml) in defined medium. Levels of Rac1, OPTN and GAPDH for loading control are assessed on 2% total lysates.

**C)** Quantification of Rac1 levels in detergent resistant membrane fractions 1, 2 and 3 as in figure 2D. Rac1 signal was first normalized to total lysate 2% (Tot). Data show fold changes compared to fraction 1 in siCtrl condition. Mean ± SD, n=3, $p^* \leq 0.05$.

**SUPPLEMENTAL FIGURE 4**

**A)** Quantification of cyclin D1 protein level in cells treated as in Figure 6B. Densitometry was normalized to value in siCtrl cells plated on 0.2kPa ECM, set to 1. Mean ± SD, n = 3 independent experiments, $p^{**} \leq 0.01$.

**B)** Immunoblots showing inhibition of cyclin D1 expression in cells treated 6 hours with the Rac1 inhibitor EHT1864 in siCtrl and siOPTN HUVECs. Immunoblots anti-OPTN and GAPDH show controls of OPTN knockdown and protein loading, respectively.

**C)** Level of cyclin D1 mRNA expression in siCtrl or siOPTN transfected cells expressing HA-Rac1Q61L. Cyclin D1 mRNA levels were assessed by RT-qPCR and normalized to siCtrl condition, set to 1. Mean ± SD, n=3, ns: non significant.

**D)** Immunoblots showing the rate of cyclin D1 expression after 15 and 30 minutes of release from protein synthesis blockage with 10 nM Cycloheximide (CHX). Immunoblots anti-OPTN and GAPDH show controls of OPTN knockdown and protein loading, respectively.

**E)** Quantification of cyclin D1 neo synthesis in cells treated as in Supplemental Figure 6C. Data show mean ± SD, n=3 independent experiments, p*≤0.05.

**SUPPLEMENTAL FIGURE 5**

**A)** Immunoblots showing OPTN depletion in transfected cells shown in Figure 5A and Supplemental figure 5B. GAPDH was used as a loading control.

**B)** Confocal sections showing enhanced formation of Zyxin-positive positive FA in siOPTN transfected HUVECs plated 2 hours on fibronectin in defined medium.

**C**) Number and mean area of FA structures positive for Integrin alpha5 in siCtrl or siOPTN transfected HUVEC were quantified. Results are expressed as fold change to siCtrl condition, set to 1. n = 30 SiCtrl cells and 27 SiOPTN cells, p*≤0.01, p**≤0.001.

**D)** Adhesion of control or Flag-OPTN expressing HUVECs on 1.25, 2.5 and 5 µg/ml of fibronectin was assessed upon short-term plating of cells in triplicate in 96 well plates. Graphs represent the extent of adhesion to fibronectin as function of absorbance upon elution of the crystal violet dye. Mean ± SD, n=3 independent experiments, p**≤0.01, p***≤0.001. The immunoblots show Flag-OPTN expression in transfected cells shown in Supplemental figure 5D and 5E. GAPDH was used as a loading control.

**E)** Confocal sections showing reduced formation of integrin alpha-5-positive (ITGA5) and paxillin-positive Focal Adhesions (arrows heads) in control or Flag-OPTN expressing HUVECs. Arrows indicate peripheral Focal Complexes. Bar = 10 [illegible]m. The square defines the region shown with merged labeling at x3.5 zoom.

**SUPPLEMENTAL FIGURE 6**

**A)** On the left: Quantification of Ec bound to cells transfected by siCtrl or siOPTN. Data represent the absolute number of CFU/mL (Y axis starts at $10^6$). Mean ± SEM of five independent experiments and 3 replicates per condition. P value: non significant. On the right: One representative immunoblot for verification of OPTN knock down. GAPDH is used as loading control.

**B)** Representative immunoblot showing transfection of the Talin-head plasmid fused to GFP compared to the empty vector (pGFP) in cells treated as in figure 6E. GAPDH is used as loading control.

**SUPPLEMENTAL FIGURE 7**

**A)** Representative spatial distribution of traction force vectors (yellow arrows) in HUVEC transfected with siCtrl or siOPTN (outlined in white) and adhered 3h to micropillar substrates coated with fluorescent fibronectin (red). White scale bar, 10 µm. Yellow scale bar = 10 nN.

**B)** Total traction force exerted by each siControl and siOPTN cell. Total force was calculated by adding the magnitude of all the force vectors in each cell. Forces are significantly lower for OPTN knocked down cells (n=32 siCtrl and n=33 siOPTN cells, p***≤0.001).

**C)** Total contractile energy stored by a cell in the substrate. The total contractile energy was obtained by adding the strain energy of all the micropillars in each cell (n=32 siCtrl and 33 SiOPTN cells, p***≤0.001).

**SUPPLEMENTARY TABLE S1 and S2**

List of quantified proteins in HUVEC cells cultured on ECM of different stiffness (n=1,804) in the non-treated (NT, Sup-table S1) or CNF1-treated condition (CNF, Sup-table S2). Pairwise statistical tests were performed to reveal proteins that were significantly regulated under different substrate rigidity conditions. Columns from left to right contain: an indication whether the protein is significantly up- or downregulated, -log10 (adjusted p-value/FDR), log2 fold change ratio, Uniprot accession, protein name, gene name and identification score. Proteins are ordered by increasing FDR value for the 50kPa to 1kPa comparison test. Information in grey refers to protein groups that contain common proteomics experiments contaminant proteins.

**SUPPLEMENTARY TABLE S3.**

List of proteins Z-scored LFQ intensities derived from the protein heatmap obtained after statistical testing and hierarchical clustering  (n=1,076). Proteins are ordered according to heatmap on Figure2A. Pairwise statistical tests were performed to reveal proteins that were significantly regulated under different ECM rigidity conditions. Columns from left to right contain: Uniprot accession, protein name, gene name, cluster number and zscore per sample (empty cells represent undetected proteins). Information in grey refers to protein groups that contain common proteomics experiments contaminant proteins.

**SUPPLEMENTARY TABLE S4**

Functional pathway enrichment analysis using DAVID. List of upregulated KEGG Pathways and Keywords for the Non treated (NT) and CNF1 (CNF) conditions. Table contains number of proteins per term (count column), P-value, list of genes and statistical tests (Bonferroni, Benjamini, FDR)

Supplemental FIGURE 1

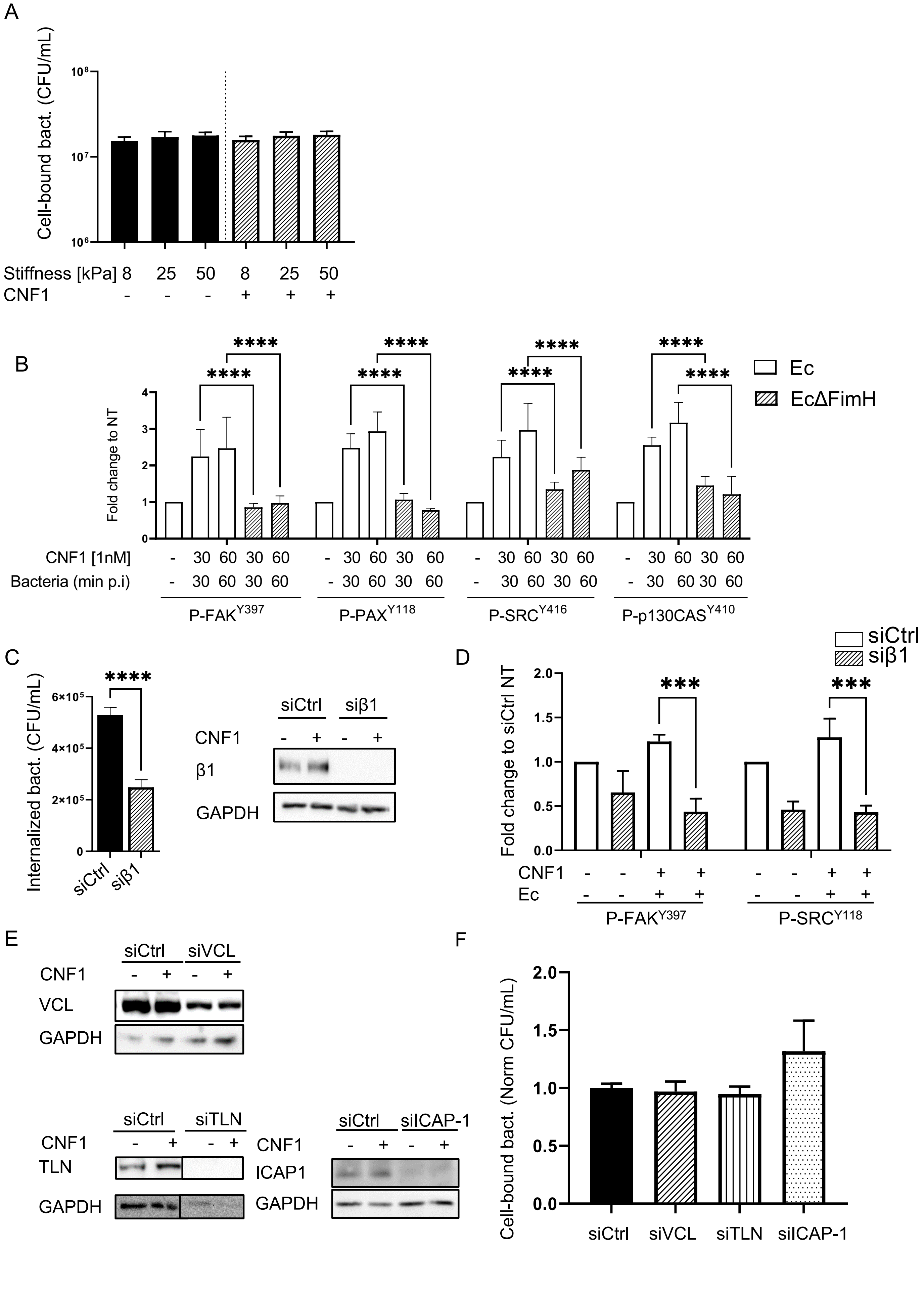

Supplemental FIGURE 2

A

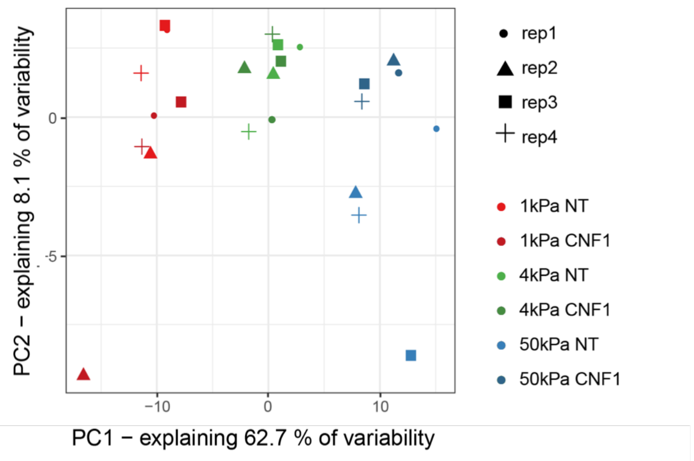


B

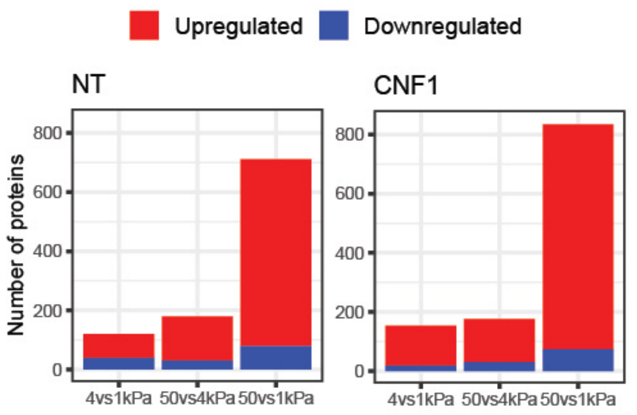

Supplemental FIGURE 3

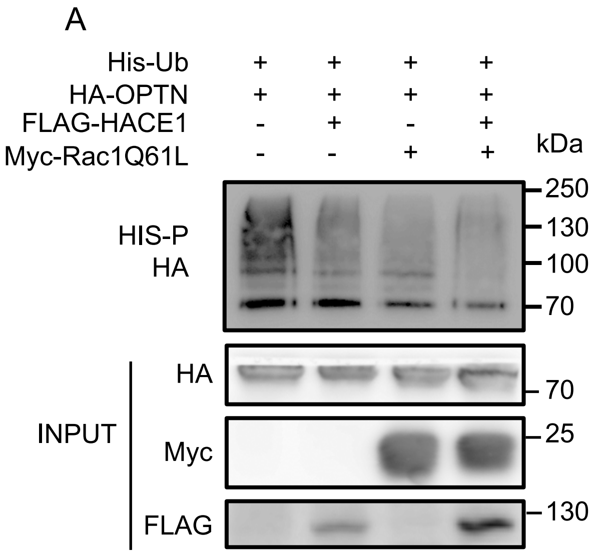


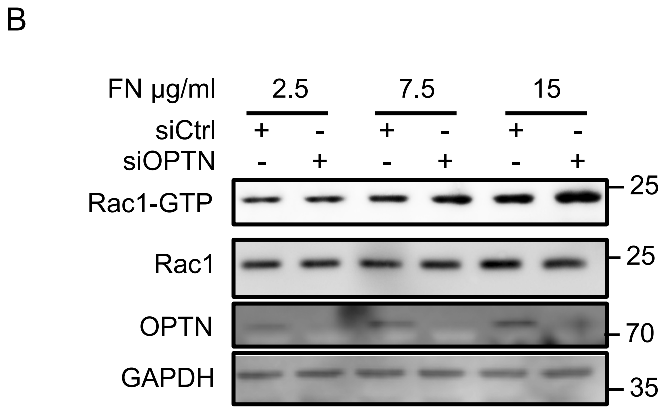


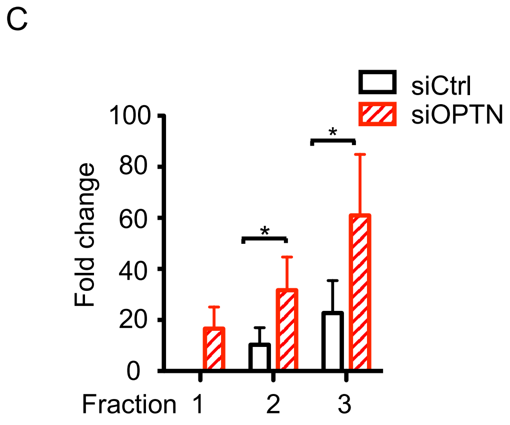

Supplemental FIGURE 4

A

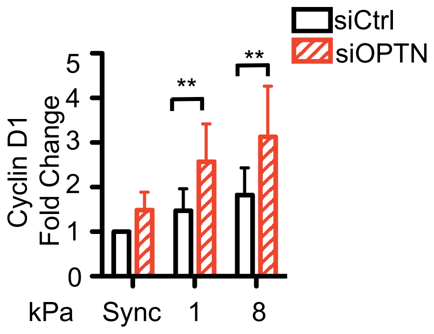


B

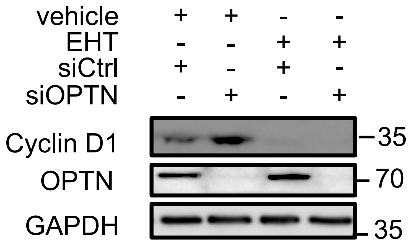


C

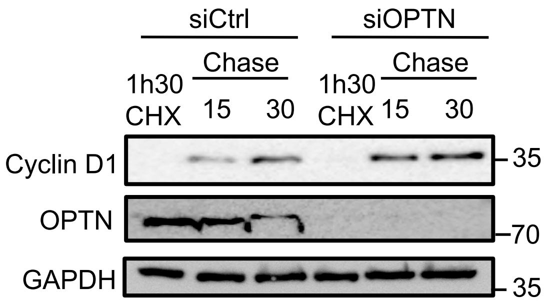


D

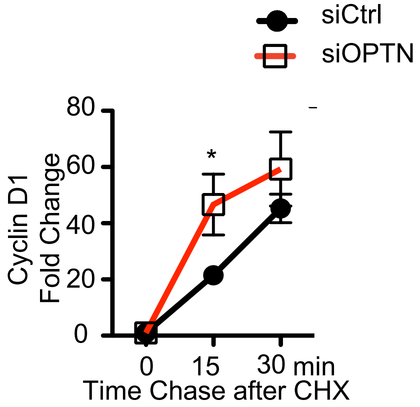

Supplemental FIGURE 5

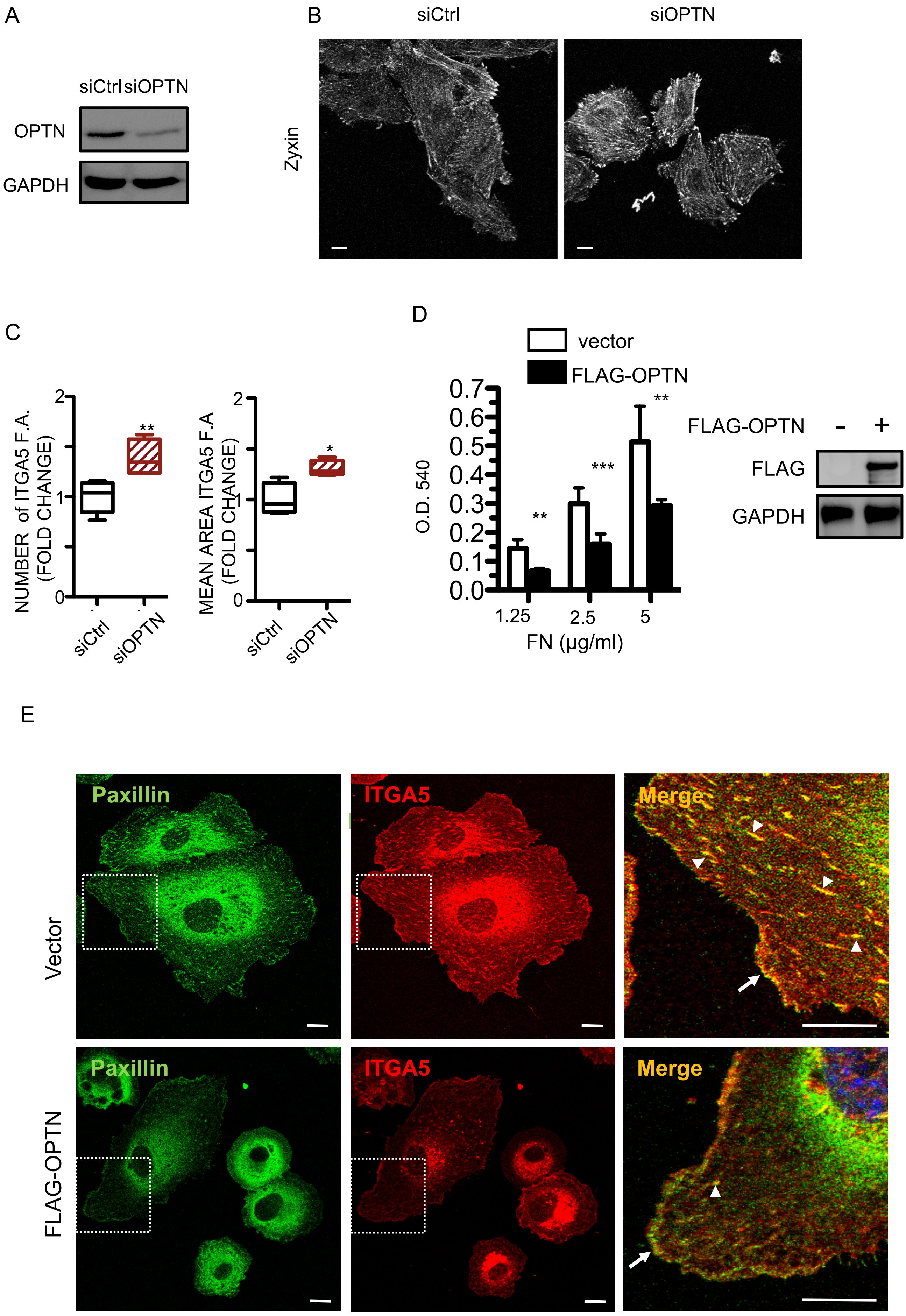

Supplemental FIGURE 6

A

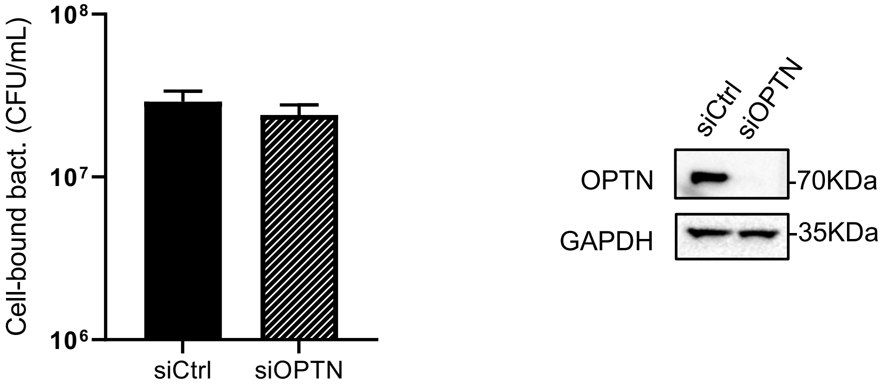


B

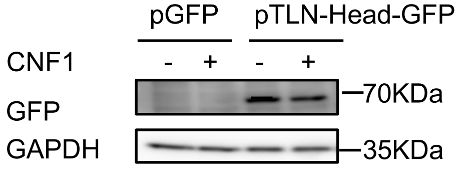

Supplemental FIGURE 7

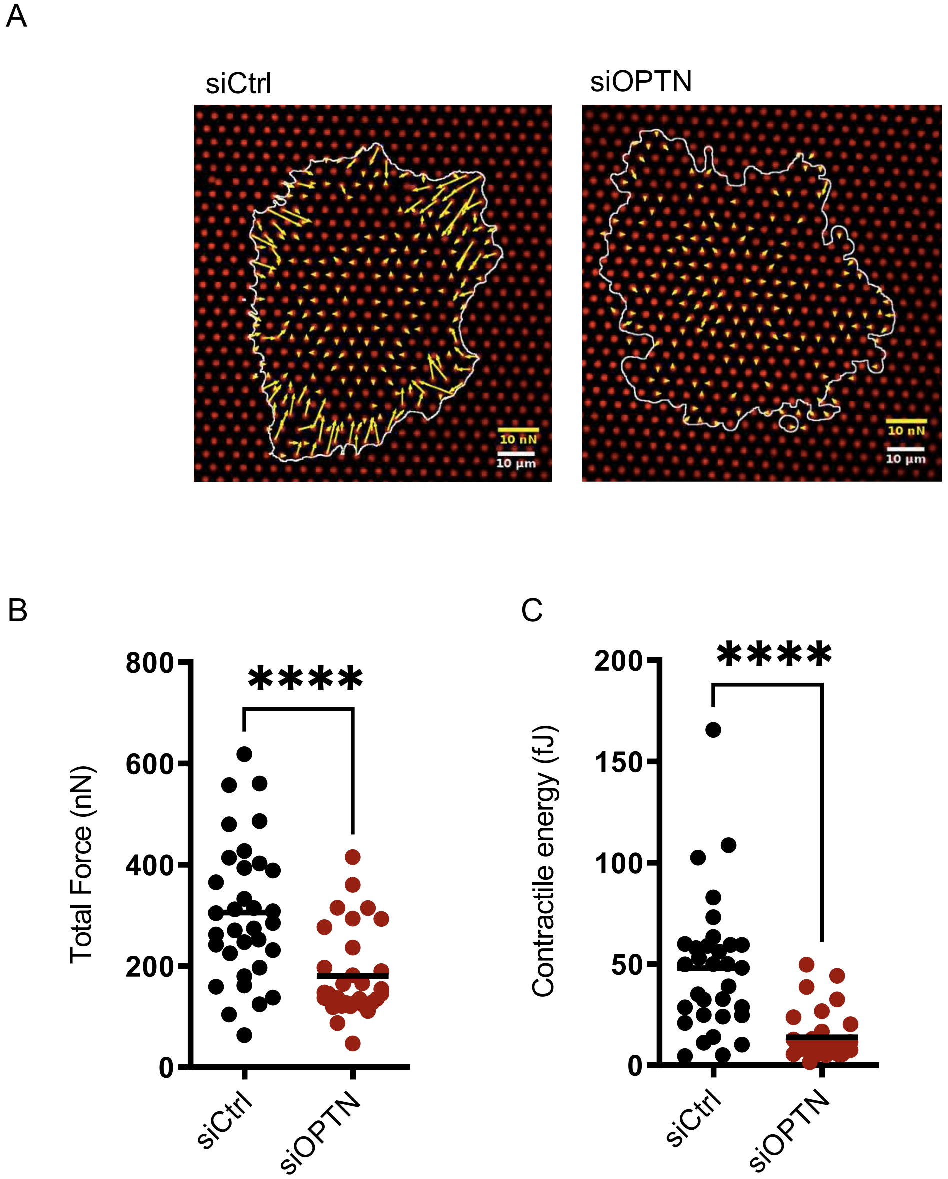
